# TorchLens: A Python package for extracting and visualizing hidden activations of PyTorch models

**DOI:** 10.1101/2023.03.16.532916

**Authors:** JohnMark Taylor, Nikolaus Kriegeskorte

## Abstract

Deep neural network models (DNNs) are essential to modern AI and provide powerful models of information processing in biological neural networks. Researchers in both neuroscience and engineering are pursuing a better understanding of the internal representations and operations that undergird the successes and failures of DNNs. Neuroscientists additionally evaluate DNNs as models of brain computation by comparing their internal representations to those found in brains. It is therefore essential to have a method to easily and exhaustively extract and characterize the results of the internal operations of any DNN. Many models are implemented in PyTorch, the leading framework for building DNN models. Here we introduce *TorchLens*, a new open-source Python package for extracting and characterizing hidden-layer activations in PyTorch models. Uniquely among existing approaches to this problem, *TorchLens* has the following features: (1) it exhaustively extracts the results of all intermediate operations, not just those associated with PyTorch module objects, yielding a full record of every step in the model’s computational graph, (2) it provides an intuitive visualization of the model’s complete computational graph along with metadata about each computational step in a model’s forward pass for further analysis, (3) it contains a built-in validation procedure to algorithmically verify the accuracy of all saved hidden-layer activations, and (4) the approach it uses can be automatically applied to any PyTorch model with no modifications, including models with conditional (if-then) logic in their forward pass, recurrent models, branching models where layer outputs are fed into multiple subsequent layers in parallel, and models with internally generated tensors (e.g., injections of noise). Furthermore, using *TorchLens* requires minimal additional code, making it easy to incorporate into existing pipelines for model development and analysis, and useful as a pedagogical aid when teaching deep learning concepts. We hope this contribution will help researchers in AI and neuroscience understand the internal representations of DNNs.

## Introduction

Deep neural network models (DNNs) have emerged as the dominant class of AI models for performing many tasks, and as promising, albeit debated, candidate models for how the brain functions (Khaligh-Razavi & Kriegeskorte, 2014; Kriegeskorte, 2015; Xu & Vaziri-Pashkam, 2021; Yamins et al., 2014). A DNN implements a series of operations that transform inputs into outputs. For many practical applications it is not necessary to examine the results of intermediate operations in DNNs. However, probing the intermediate steps can help us understand the computations used by a particular model to transform inputs into outputs, illuminating the reasons for its successes and failures and informing the development of better models. In addition, neuroscientists seek to evaluate how well DNNs model brain function. This requires comparing not just their performance on task benchmarks, but also the degree to which their intermediate representations match those observed in brains.

Advancing the practical goal of improving DNN task performance and the scientific goal of comparing DNNs to brains often involves comparing many different models, making it desirable to have convenient methods at hand for extracting the results of these intermediate operations and understanding their role in the information processing of each network. Ideally, such a feature extraction method should have the following qualities. First, it should easily work for *any* DNN without requiring model-specific tinkering by the user, such as editing of a model’s code, since this approach can become time-consuming and error-prone when examining many models. Second, it should be able to extract the results of *all* internal computations within a model, rather than just a subset. Third, it should render transparent—for instance, through visualization or through intuitive metadata—where a given layer fits within the broader information flow of a network. This is essential because the function of a layer depends on the other operations performed in a network, and models often include architectural complexities such as branching or recurrence. Finally, given the infinite space of possible DNN models that can be constructed, and the rapid pace of model development, it is desirable to have an automated method for validating the correctness of the extracted activations for novel models.

None of the currently existing methods of feature extraction fulfills all the above desiderata (Table 1). We focus here on models implemented in PyTorch (Paszke et al., 2017, 2019), one of the most popular libraries for implementing DNNs for both academic and industrial applications. One approach to feature extraction is to simply edit a model’s code to return the results of any intermediate operation. This approach can be time-consuming and error-prone, especially when we have to edit the code for many models. Another option is to use PyTorch’s built-in functionality to attach “forward hooks” that return the results of PyTorch *module* data structures. PyTorch modules provide a convenient way of flexibly organizing model code and ensuring that a training optimizer can access a model’s parameters. The user can additionally attach “forward hooks”: functions linked to a module that are executed when a tensor enters or leaves that module. This is a popular approach to PyTorch feature extraction because it requires no direct modification of a model’s code. However, it comes with several limitations. First, there is no universal naming convention for characterizing the modular organization of DNNs. A user examining a collection of models must become acquainted with the nomenclature of all of them. Second, and more importantly, PyTorch operations not associated with a module cannot be extracted using this approach. These include many elementary operations (e.g., tensor addition, cosine) that have no module equivalent (unless the model designer chooses to encode them as such), and operations that the model designer has implemented as functions rather than modules. For example, a ReLU operation can be implemented either as a module object (where its activations can be extracted using the forward-hooking approach), or simply as a function call (where its values cannot be extracted in this way). Thus, the ability to extract activations from a given intermediate DNN operation in PyTorch with the forward hook approach often depends on computationally irrelevant programming choices made by the model designer, which can complicate the construction of an analysis pipeline that examines many models. Put differently, while PyTorch modules are indispensable units of code organization, they do not directly map onto the elementary computational “units” of DNNs—single operations on tensors—making them suboptimal for the purpose of exhaustively characterizing the internal operations of a network.

**Table 1.** Comparison of different PyTorch feature extraction methods

|  | Ease of Use | Model Coverage | Layer Coverage | Ways of illuminating module structure |
| --- | --- | --- | --- | --- |
| **TorchLens** | Easy | All models | All layers | Visualization, layer metadata |
| **Manually Editing Model Code** | Hard | All models | All layers | None |
| **Attaching Forward Hooks to Models** | Medium | Effectively all | Only module outputs | None |
| **ThingsVision** | Easy | Pre-specified model library | Only module outputs | None |
| **Torchvision feature_extraction** | Medium | Models with static computational graph | All layers | None |
| **Surgeon-PyTorch** | Easy | Models with static computational graph | All layers | None |

In addition to these manual approaches for extracting intermediate activations from DNNs, three open-source packages now exist for facilitating this process. *ThingsVision* (Muttenthaler & Hebart, 2021) is a Python package with user-friendly functionality for loading pre-existing DNNs, extracting their activations either to common image datasets or to user-defined datasets, and performing various common analyses on the extracted features, such as representational similarity analysis (Kriegeskorte et al., 2008). While it can extract activations from a large and convenient library of pre-existing models, it does not automatically work for arbitrary new models (e.g., user-designed models), and can only extract the outputs of PyTorch modules, not every operation in the model. Another approach is the feature_extraction module provided in *Torchvision* (Marcel & Rodriguez, 2010), which works on novel models (not just a predefined library), and can extract the results of non-module operations, but does not support models with dynamic control flow, where the computational graph of a network can vary across forward passes (Looks et al., 2017; Yu et al., 2018). Examples of such models include recurrent neural networks that execute a varying number of loops until a given criterion is reached (e.g., Spoerer et al., 2020), reinforcement learning models where the agent’s behavior can vary stochastically or based on model inputs (Mnih et al., 2015), and graph convolutional networks where the computational graph can vary across model inputs (e.g., Kearnes et al., 2016). A third package is *Surgeon-PyTorch* (Schneider, 2022). Conveniently, this package not only extracts intermediate activations from both module and non-module operations, but can also isolate subcomponents of models such that they can be separately trained. However, this approach also does not work for models with dynamic control flow.

Here, we introduce a new Python package, *TorchLens*, that overcomes the limitations of these approaches: it works for arbitrary PyTorch models (not just predefined models or models with a static computational graph) and can extract the results of all intermediate operations (not just module outputs), requiring just a single line of code from the user. The ability to extract information about every operation in the model’s computational graph comes with three substantial further benefits. First, it enables the model’s computational graph to be automatically visualized, revealing the structure of the model and where a given layer is located within it. The computational graph provides a valuable aid for understanding a model’s architecture without manually inspecting its code. This visual representation of the model can also help with layer selection, especially for architectures containing complexities such as branching or recurrence, where indexing a layer by an ordinal position (e.g., conv9) may not clearly convey where a layer is situated in the network. Second, tracing all intermediate operations of a model enables producing a full characterization of the model’s computational graph—encompassing the child-parent relationships between layers, the functions executed by each layer, and so on—such that the user can programmatically select layers meeting certain criteria (e.g., all pooling layers that follow a ReLU layer), perform analyses that might draw on graph-theoretic properties of the network (e.g., the number of convolutional layers intervening between two target layers of interest), and characterize other useful properties of the network, such as the execution time required for different kinds of layers. Finally, extracting the results of *all* intermediate operations enables an automated approach for verifying the correctness of saved activations without manual troubleshooting. This verification method re-executes the functions associated with each layer using the saved activations of the inputs to that layer and checks that the resulting downstream model outputs match the ground-truth outputs. Given the large space of possible DNN models, and the fact that feature extraction bugs can easily yield “silent errors” (for instance, if a tensor of the correct dimensions but the wrong values is saved), verification ensures that feature activations are being correctly extracted for novel models that may have complex architectures.

## User interface

### Example workflow

Figure 1 depicts a simple “minimal use-case” example of how to use *TorchLens*. The user simply passes a model and input into the main function of *TorchLens*, get_model_activations, and receives a ModelHistory object that encodes information both about the entire model and about each individual layer, including the saved activations for each layer. Layer information is retrieved simply by indexing the ModelHistory object with any valid layer label. If specified, the same function call also automatically generates a visualization of the model’s computational graph (Figure 2).

**Figure 1:**
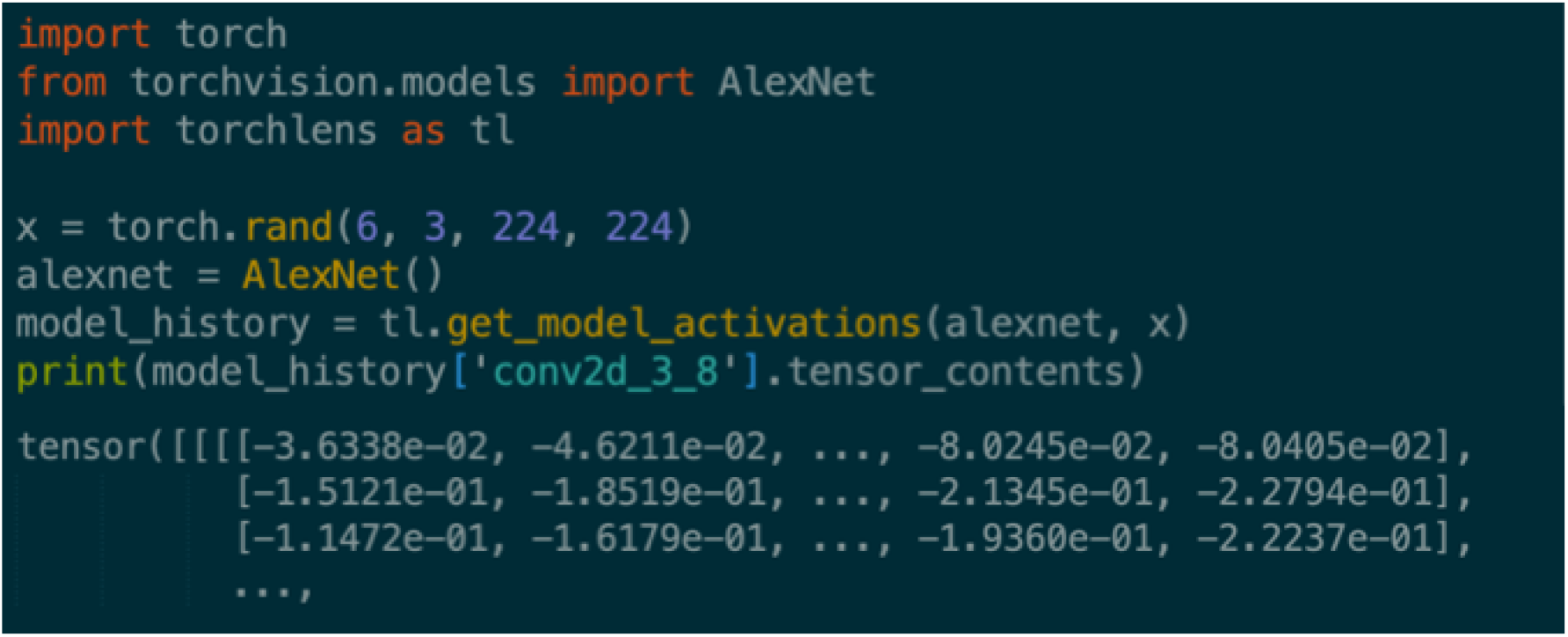
Example workflow for TorchLens. The user simply passes in the model as-is and an example input, and TorchLens returns a ModelHistory object containing information about the model and its layers, including saved activations.

**Figure 2:**
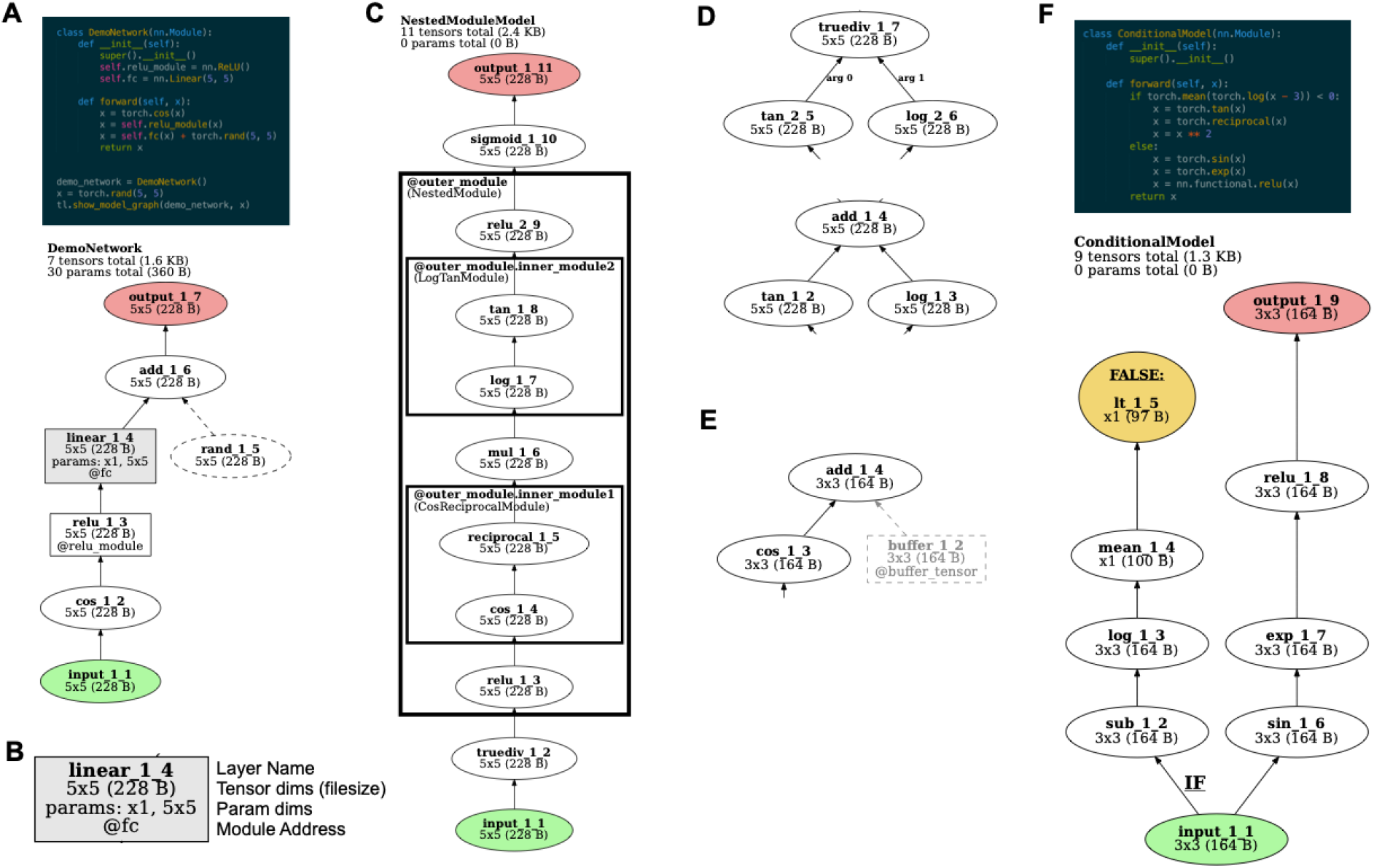
*TorchLens* visualization features. **A.** Code and visualization for a simple example model. Graph nodes correspond to tensor operations, edges correspond to parent-child relationships between operations (i.e., that the output of one operation is the input to another). Input operations are shown in green and output operations are shown in red. Operations in a module are shown as a box; other operations are shown as ellipses. Nodes for operations that create a new tensor inside of the model (e.g., from torch.rand) are rendered with dashed lines. The visualization also shows the name of the model in the top left, the number of tensors computed in the model and their total file size, and the number of trainable parameters in the model along with their total file size. **B.** The node for each layer contains the label of the layer, the shape and file size of the tensor returned by that layer, the shape of any trainable parameter tensors involved in that layer, and the module address for that layer (i.e., how it would be indexed from the Python model object) if relevant. **C**. For models containing (possibly nested) modules in which multiple operations are performed, these modules are rendered as boxes surrounding the nodes for those operations, along with a label giving both the address of that module within the model, and the class of the module. **D.** If an operation is non-commutative and takes multiple inputs (e.g., division), these inputs are marked with their argument position in the function call (e.g., 0 for the numerator and 1 for the denominator for the PyTorch division function). **E.** “Buffer” tensors that are stored along with the model, but do not contain trainable parameters, are rendered in light gray. **F.** For models containing dynamic control flow with if-then branching, the operation returning the final Boolean value in evaluating the “if” statement is marked in yellow, and the full set of operations involved in evaluating the “if” statement is labeled with a boldfaced “IF”.

### Summary of user-facing functions

*TorchLens* consists of a small handful of user-facing functions, which take as input any arbitrary PyTorch model and the input to that model. For ease of use, the main functionality of *Torchlens* is provided via the “core” function get_model_activations. This function extracts the activations from either all layers of a model or a desired subset along with metadata about the model and layers, returns a Python object of a class, ModelHistory, that the user can easily index to retrieve this information, and optionally produces a visualization of the model’s computational graph (Figure 2). The next two functions provide subsets of the functionality of get_model_activations. The function get_model_structure extracts the metadata about a model and its layers without saving any activations. This function is useful in cases where the user wishes to analyze the computational structure of a model without saving the activations, or where the goal is to programmatically select which layers to save in a subsequent call to get_model_activations (e.g., all ReLU layers that follow a convolution layer). Another function, show_model_graph, visualizes the model graph without saving any activations. Finally, the function validate_saved_activations runs the model on a specified input, and executes a procedure, described below, that validates the accuracy of the saved activations. All *TorchLens* functions allow the user to supply a random seed as an argument to ensure reproducible results for models with stochastic components, such as dropout layers or random noise.

### Layer nomenclature

Since there exists no universal nomenclature for referring to DNN layers, and since PyTorch models often have a nested modular design where each module is assigned a name by the model’s designer, *TorchLens* adopts the convention of using a “default” layer naming scheme that is common across all models, while also allowing the user to select the results of intermediate layers based on the module of which they are the output—helpful in cases where the model’s designer has demarcated meaningful computational “chunks” in the model using modules, or where the user is already familiar with the modular structure of a given network.

The default name of a layer follows the format {layer_type}_{layer_type_num}_{layer_total_num}; for example, “relu_3_6” refers to the third ReLU layer in the network, and the sixth layer overall. While providing the layer’s ordinal position both among layers of the same type and among all layers in the network is informationally redundant, providing both of these numbers is helpful in getting a quick sense of where the layer falls within the network. Given the common convention of referring to a layer with the ordinal position among layers of the same type (e.g., “conv2” is the second convolutional layer), we adopt the practice that while the full layer label is always provided in outputs, the user can also refer to layers with a truncated label that only specifies the first index (e.g., just “relu_3” instead of “relu_3_6”). In cases where a model has a branching structure (i.e., where a layer may have multiple output layers), such that layers do not have a unique ordinal position, numbering is based on the programmatic execution order of the layers in the code, which guarantees that a given layer always has a lower positional index than any downstream layers (i.e., they are topologically sorted). In case a model is recurrent and a layer contains multiple passes, the specific pass of a layer is indicated with a colon; for instance, conv2d_3_7:2 is the second pass of the third convolution layer (and seventh layer overall). The logic of assigning operations to the “same” layer is described in the section *Handling of recurrent networks*.

In addition to this default naming convention, which provides a way of referring to layers that is common across models, layers are also labeled based on the module for which they are an output. To provide a specific example, the PyTorch implementation of AlexNet (Krizhevsky et al., 2012) consists of a nested design with two top-level modules: a “features” module containing the convolutional layers, and a “classifier” module with the fully-connected layers, each of which itself contains a module for each of the individual operations in that block. The second convolutional layer of AlexNet is the fourth submodule of the “features” module, so the “module name” of that layer would be features.3 (following Python’s zero-indexing convention). Under the default naming convention of *TorchLens*, the same layer would be indexed as conv_2_4 or conv_2. Layers can be fetched with their “module address” at any level of nesting: for instance, the output of the third pooling layer in AlexNet is the output of module “features.12”, but also of the higher-level module “features” (of which “features.12” is the last submodule).

### The ModelHistory object: a full characterization of the model’s forward pass

The core user-facing data structure used by *TorchLens* (returned by get_model_activations and get_model_structure) is the ModelHistory object, which contains both metadata about the entire model, and information about each individual layer, including both metadata and the saved activations for that layer if specified by the user. Broadly, the provided metadata includes information about the overall structure of the network’s computational graph, the tensor returned by a given layer, the function executed by each layer (including any trainable model parameters associated with that function, such as convolution weights), and the parent-child relationships between layers along with further graph metadata (e.g., the number of layers separating a given layer from the input or output). This metadata also includes useful performance-relevant data for a network: for instance, it includes the execution time of each layer, the memory size of the tensor returned by each layer, and the memory size of any trainable model parameters associated with a layer, facilitating profiling of the execution time and memory usage of a model. Supplementary Table 1 provides a complete list of the metadata provided by *TorchLens*. Simply printing the ModelHistory object provides an overall summary of the model, with further metadata available by indexing the attributes of the object. Data about each layer can be retrieved by indexing ModelHistory, using any of several valid labels for a layer: the “default name” of a layer, such as conv2d_3_7 or conv2d_3, the “module name” of a layer, such as “features.8”, or the overall ordinal position of the layer in the network, following the conventions of Python list indexing (e.g., model_history[2] is the third layer, and model_history[-5] is the fifth-to-last layer. If only a subset of layers is saved, numerical indexing is performed with respect to the saved layers only.

### Visualization

In addition to returning the ModelHistory data structure with information about the forward pass, *TorchLens* can also automatically produce a visualization of the model’s computational graph (Figure 2). The visualization enables the user to easily understand the structure of the model and a given layer’s place within that structure, aiding in both the overall understanding of a model, and in the selection of particular layers from which to extract activations. *TorchLens* uses the graphviz library (Ellson et al., 2004) to programmatically draw the computational graph without requiring the user to manually position the nodes.

The visualization produced by *TorchLens* is designed to make the overall organization of the network and the important properties of individual layers visually salient. Figure 2 depicts the visual code used by *TorchLens*. Each node is a particular layer, and the directed edges (arrows) show parent-child relationships between layers (i.e., the layers whose output feeds into subsequent layers). Input layers are shown in green, output layers are shown in red, and layers whose functions have trainable parameters are shown in gray. Elliptical nodes are single operations that are not associated with a module, whereas rectangular nodes are associated with a “bottom-level” module associated with a single operation; for instance, a ReLU operation implemented as a function call would be shown as an ellipse, and a ReLU operation connected with a module is shown as a rectangle. In addition to these “bottom-level” modules that only execute a single tensor operation, PyTorch contains “higher-level” modules that encompass multiple operations, which are shown as (potentially nested) rectangles surrounding the nodes corresponding to layers contained within those modules (Figure 2C).

The visualization includes metadata both about the overall model, and about each individual layer. For the overall model, the visualization shows the name of the model class, the number of tensor operations in the model along with the total memory size of the resulting output tensors, and the number of trainable parameters in the model, along with the total memory size of these parameters (Figure 2B). The node corresponding to each layer in the model is marked with 1) the layer’s “default” name (e.g., conv2d_2_4 for the second convolution layer, and fourth layer overall), 2) the dimensionality and memory size of the output tensor (useful for choosing which layers for which to save activations), 3) the shape of any associated parameter tensors for that layer (if applicable), and 4) the “module address” of that layer if it associated with a module. The box outlines corresponding to higher-level modules are also marked with this address.

Finally, TorchLens indicates several other kinds of information to remove ambiguity about the structure of models that include less common “special case” components. Some PyTorch models contain internally-generated tensors that are not computed from the model’s input, but contribute to the model’s computations (e.g., if randomly generated noise is injected at some stage of processing via a function like torch.rand). These operations are indicated with dashed lines, intended to make it easy to visually distinguish them from layers whose inputs originate from model inputs, such that the user can visually trace the path from input to output (Figure 2A). Some PyTorch models contain hard-coded “buffer” tensors stored along with a model that are not trainable parameters; these are rendered in greyscale (Figure 2E). In cases where a layer performs a non-commutative operation (e.g., division) on the outputs of multiple parent layers (Figure 2D), the function argument number of each edge is explicitly marked (e.g., “arg 0” for the incoming layer whose output tensor serves as the numerator, and “arg 1” for the denominator layer) for positional arguments, and the name of the argument is marked for keyword arguments. In cases where a network contains conditional control flow, where an if-then statement is evaluated on a tensor to yield a single Boolean value (e.g., to perform different computations based on the value of the model’s input), *TorchLens* marks the node yielding this single Boolean value in yellow, infers the beginning of the set of computations involved in computing this Boolean value, and marks the beginning of the branch of the computational graph involved in computing this Boolean value with a bold “IF” label (Figure 2F).

We include a [publicly available collection](https://drive.google.com/drive/u/0/folders/1BsM6WPf3eB79-CRNgZejMxjg38rN6VCb) of visualizations for over 800 commonly used image, video, audio, and language models, demonstrating the generality of this automated visualization approach.

### Handling of recurrent operations

Since many DNN models contain recurrent (feedback) processing, *TorchLens* has built-in procedures that automatically identify and demarcate recurrent layers, using the convention of marking passes of a layer with a colon as noted above (e.g., conv2d_2_4:3 for the third pass through this layer).

An important conceptual question to address is what counts as a “recurrent layer”. At a ground-truth level, a DNN is simply a series of tensor operations, some of which have trainable parameters associated with them. A complete description can be given of a network in this way without invoking the notions of “feedforward” and “recurrent” processing. Nonetheless, distinguishing feedforward and recurrent operations can be helpful in clarifying the computational structure of a network. We stipulate that in a “feedforward” network, each layer executes one operation, but in “recurrent” networks a layer may execute its associated operation multiple times during a forward pass. Thus, identifying “recurrent” layers corresponds to finding sets of operations in the computational graph that can be usefully regarded as belonging to “the same layer”. The most obviously useful operations to treat in this way are ones in which functions with the same trainable parameters are executed multiple times during a forward pass. For example, if a convolution operation with a given set of weights is executed three times over the course of a network, these operations should be regarded as being associated with the “same” layer. However, other operations may also be usefully regarded as “the same”. For instance, if each of the three aforementioned convolutional layers is followed by an identical ReLU layer and a pooling layer, it is also sensible to group together these three ReLU operations as belonging to “the same” layer, and likewise for the three pooling layers. Finally, if a repeated and contiguous series of operations with no stored parameters is executed (e.g., XYZXYZXYZ, where X, Y, and Z are different functions), it is also useful to regard this as a “loop” in the graph, and to mark corresponding operations in the loop as the same layer.

*TorchLens* thus adopts the following conventions (illustrated in Figure 3) for marking layers as “recurrent”:

1. operations with associated trainable parameters that are executed multiple times in the forward pass are always marked as passes of the same layer,
2. all repeated operations adjacent to these repeated-parameter layers are marked as passes of the same layer, irrespective of whether the “chunk” of layers associated with a pass of a parameter layer ends up being adjacent to a chunk associated with another pass of that parameter layer, and
3. if there are repeated, adjacent sequences of operations without trainable parameters, then corresponding layers in these sequences are marked as the same, but not if these sequences are separate (e.g., XYZXYZXYZ is recurrent, but XYZAXYZBXYZ is not, since the XYZ operations are not adjacent).

**Figure 3:**
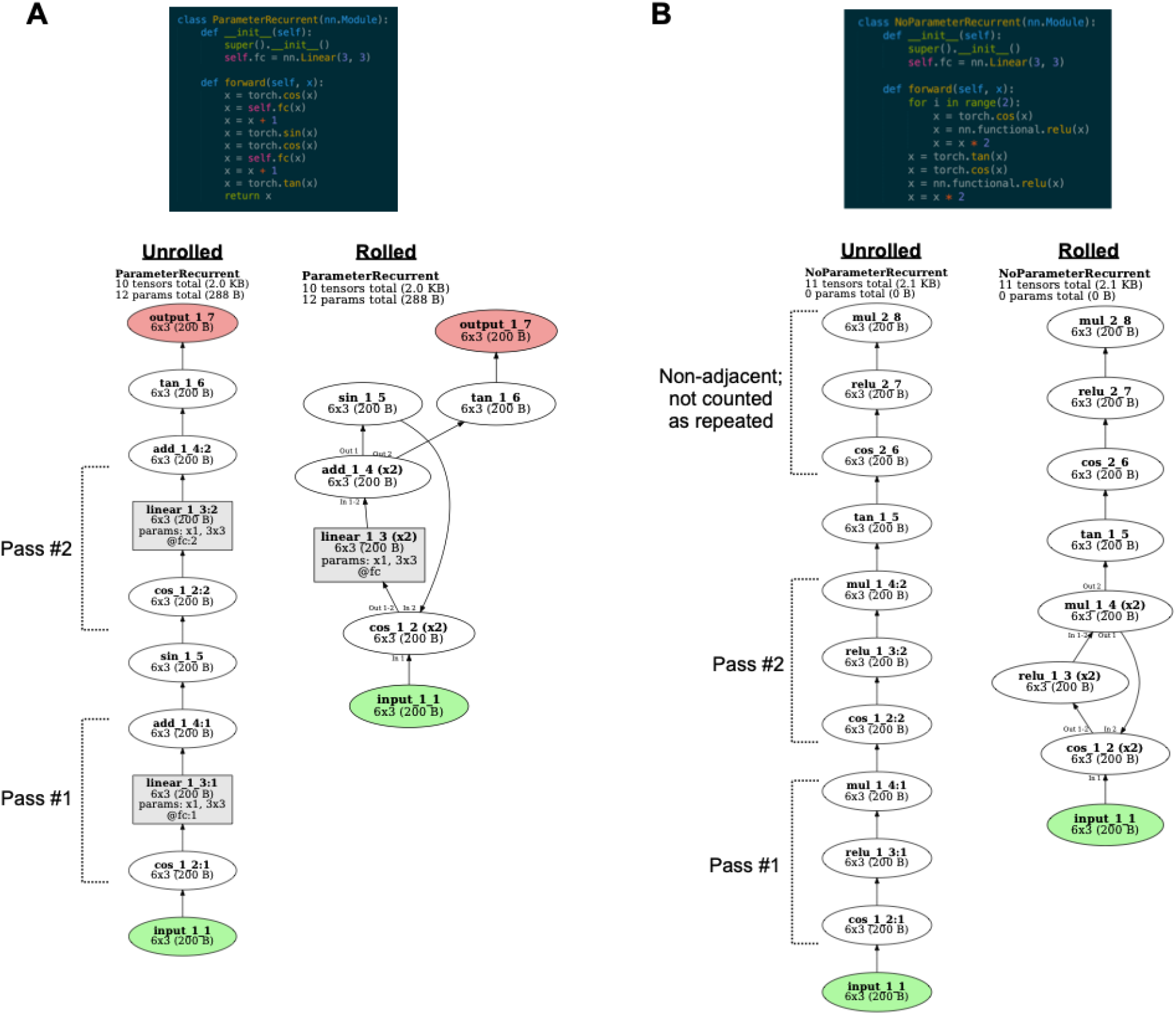
Definition and visualization of recurrent layers. **A.** Operations are considered as passes of the same layer if the same parameters are repeated (e.g., here the same fully-connected operation parameters are applied twice), or if there are operations that are both identical to each other, and contiguous to repeated-parameter layers (e.g., here the same cosine operation is applied prior to both passes of the fully-connected layer, and the same addition operation is applied after both passes). For the unrolled visualization, each pass of a layer is rendered with a separate node, with the pass number marked with a colon (e.g., linear_1_3:2 for the second pass of layer linear_1_3). For the rolled visualization, all passes of a layer are marked with the same node, the number of passes for that layer is given in parentheses (e.g., “(x2)”), and in cases where a layer has different input and output layers for different passes, these passes are labeled next to the incoming or outgoing arrows to the node for that layer. For instance, the layer cos_1_2 takes input from input_1_1 for its first pass, but sin_1_5 in its second pass. **B.** Operations are also considered as passes of the same layer if they do not contain trainable parameters, but repeat as part of an identical sequence. For instance, here the same sequence of cosine, ReLU, and multiplication operations is repeated twice in a row, so the two repetitions of each operation are assigned to the same layer. However, the same series of three operations occurring later (after the intervening tangent operation) are not grouped together with the earlier corresponding layers, since they are not contiguous with other occurrences of this sequence.

Note that these are simply conventions to aid in quickly understanding a model, since whether a layer is “recurrent” is to some degree a matter of definition.

For visualizing recurrent models (Figure 3), *TorchLens* also includes functionality for showing the model in both unrolled format, where separate operations of a recurrent layer are shown as different nodes, and in rolled format, where separate operations of a recurrent layer are shown as just one node (Figure 3). In the unrolled format, each node is marked with the pass of the layer associated with that node: for instance, linear_1_1:2 is the second pass of the first linear layer. In the rolled-up format, recurrent layers are marked with the number of passes they perform next to the name of the layer. For instance, “linear_1_1 (x3)” means the layer has three passes. In some cases, a layer’s inputs and output layers can vary across passes. For example, perhaps in the first two passes a layer sends feedback to the beginning of a loop of which it is a part, but in the third pass it sends activation out of the loop. *TorchLens* explicitly marks such cases when they occur by indicating the pass number on incoming and outgoing edges (e.g., “In 1” next to an incoming edge means that input comes from the associated parent layer during the first pass, and “Out 2-3” next to an outgoing edge means that output is sent to that child layer in the second and third passes).

### Validation

The approach used by *TorchLens* (described in the “Implementation” section below) can in principle be used for any PyTorch model. The space of possible DNN models is infinite because the number of ways that tensor operations can be assembled into a computational graph is unlimited. It is important to have confidence that the activations saved from these models are correct, even for novel models with complex or exotic architectures that we may not have anticipated while developing *TorchLens*. This is especially important because of the possibility of “silent errors”, where a given DNN operation (e.g., a ReLU operation) may change the values of a tensor without changing its shape, such that an error in saving the activations would be harder for the user to detect. Since manually checking the correctness of saved activations for a new model can be time-consuming and error-prone, *TorchLens* contains a built-in procedure for algorithmically validating the accuracy of saved activations. This is achieved by re-executing the function associated with each layer with the saved tensor values of the layer’s parents, and verifying that the resulting tensor matches the saved tensor for the layer. This procedure is described in more detail in the “Implementation” section. The user can execute this procedure using the validate_saved_activations function on a given model and input, which will return True if the saved activations for that input pass the validation test, and False if they do not.

So far, this procedure has been internally run on a large collection of DNNs, including 1) all image and video models in the TorchVision model zoo (Marcel & Rodriguez, 2010), including various visual transformer models, 2) over 700 models in the TIMM model zoo, 3) the CORNet class of image models inspired by the primate ventral visual pathway (Kubilius et al., 2018), 4) all models in the TorchAudio model zoo, and 5) the language models BERT (Devlin et al., 2019) and GPT2 (Radford et al., 2019). Based on these extensive tests, we believe *TorchLens* works robustly. In case any architectural “edge cases” are not currently accounted for by TorchLens, this built-in validation procedure will detect such cases and allow them to be corrected in subsequent updates to *TorchLens*.

### Example full workflow

As a way to summarize the elements of the user interface described above, we briefly describe an imagined use-case for *TorchLens:* a user has either designed their own PyTorch model or downloaded a model, and wishes to extract the activations from a selected set of layers.

#### (1) Inspect the model graph

First, to get a sense of how large the model is and which layers may be interesting to examine, they call the show_model_graph function to visualize the computational graph of a model without saving any activations. They note that the network is very large, such that saving the activations of all layers may use up a prohibitive amount of RAM, and selecting only a subset of layers to save may be desirable. They note that the model is organized into several high-level modules, each of which contains multiple recurrent feedback loops, and further note that the model contains many cases where a pooling layer follows a ReLU layer.

#### (2) Optionally use model structure to find representations of interest

They decide they wish to extract the outputs of all high-level modules in the model, the end of each pass of the recurrent loops within the modules, and the results of all pooling layers that follow a ReLU layer. To identify these ReLU-pooling layer pairs, they call get_model_structure to extract the metadata of the model, loop through each layer’s metadata, and find all pooling layers that have a ReLU layer as a parent.

#### (3) Extract activations for representations of interest

Finally, they call get_model_activations, specifying the desired layers whose activations they wish to save, and verify that the activations were accurately saved for this new model by calling validate_saved_activations for the model and input.

## Implementation

### Logging the forward pass

We now describe the internal implementation of *TorchLens*. As laid out in the introduction, we sought an approach that could automatically extract the results of any tensor operation from arbitrary PyTorch DNN models without manually editing model code. As noted earlier, currently existing approaches face various limitations: for example, they can extract the results of module operations but not the results of operations that are not linked to modules, or they only work in the case of models with a static computational graph, but not for models with a dynamic computational graph that can vary across forward passes (e.g., via if-then branching).

To overcome these limitations, we devised a dynamic tracing approach that relies upon Python’s built-in decorator functionality in order to transiently modify all built-in PyTorch functions that return a tensor. These modified functions log information related to each tensor operation. A decorator is a Python function that wraps another function with additional code that is executed whenever that function is called. To extract the results of the tensor operations in a model, we constructed a decorator that inspects the inputs and outputs of a PyTorch function call, and logs information about the function and its inputs and outputs to a Python object of a class called ModelHistory that we implemented for this purpose. Additionally, any output tensors are tagged with a “barcode” (simply an identifier attribute assigned to the tensor data structure) such that when any subsequent functions are called on that tensor, the parent layers feeding into that function can be looked up from ModelHistory, thus establishing the parent-child relationships in the computational graph. To log the results of the forward pass of a model, *TorchLens* applies this decorator to all PyTorch functions that return a tensor as output, and a forward pass is then run on the model. The model’s code remains unchanged, but the functions called within that code now are decorated with the additional logging functionality, and thus log their results to ModelHistory. Additionally, in order to log the modules in which operations occur, *TorchLens* also attaches a forward hook to each module that tracks whenever a tensor enters or leaves the module and logs this to ModelHistory. When the forward pass is completed, the logging decorators are removed from the PyTorch functions, returning them to their original definitions.

The advantage of this approach is that PyTorch contains a limited (though still substantial) set of elementary operations from which nearly all higher-level functions are constructed; thus, so long as *TorchLens* properly handles the logic of logging the results for each elementary operation, it should in principle be able to work for all models built from the elementary operations, instead of being limited to a predefined library of models. We note that this approach sometimes requires dealing with various “special case” functions. For example, some functions change their input tensors in place and either return nothing or return the same tensor object (without creating a new tensor), and some return a list or tuple of tensors rather than a single tensor. Each of these cases must be specifically dealt with, and it is possible that new elementary operations with unique programming logic will be added to PyTorch with further updates. However, PyTorch’s elementary operations will always constitute a finite set, and it should be straightforward to address them as they arise with further updates to *TorchLens*. Furthermore, since *TorchLens* also contains a built-in validation procedure (implementation described below), it can detect these special cases when they arise. Thus, the approach used by *TorchLens* should be both broadly generalizable to any PyTorch model, and easily adapted to any potential further additions to PyTorch’s set of elementary operations.

### Post-Processing the Model’s Computational Graph

Following the forward pass, several post-processing operations are applied to the stored computational graph. First, “orphan operations” that neither arise from the model’s input nor contribute to the model’s output are trimmed from the graph. Second, the number of layers separating each layer from the model’s input and output layers is computed. Third, *TorchLens* attempts to infer cases where the model contains conditional branching via if-then statements. To do this, it identifies and marks operations whose output is a tensor consisting of a single Boolean value that is not subsequently fed into any further operations, which is a plausible candidate for an operation evaluated in an “if” block. While this is not a bulletproof heuristic, there are few cases in which a coder would compute a childless Boolean tensor in this way that is not part of an “if” block. Additionally, *TorchLens* traces backwards from the layer whose output is this Boolean tensor until it finds a layer that is an ancestor of the model’s output, and marks this transition point as the beginning of an “if” branch. For example, if an “if” statement consists of “if torch.mean(x-5) > 3”, the “greater than” operation would be the childless node that returns a single Boolean value, and the subtraction operation would be the first operation that branches off from the input and leads into the series of calculations involved in evaluating this if-statement (as in Figure 2F). Tagging if-statements in this way enables them to be marked on the visualization of the computational graph produced by *TorchLens*, clarifying the control flow of the forward pass.

Fourth, *TorchLens* corrects the module containment information (i.e., which of the possibly-nested modules the operation occurs in) for operations performed on internally generated tensors—for instance, via functions such as torch.ones, or torch.rand. Since these tensors can be generated inside a module, the module in which they are generated cannot be inferred on the fly by tracking the modules entered by any ancestor tensors. To correct this, *TorchLens* recursively traces the children of these layers until it finds a layer that descends from the model input—which will have accurate module containment information—and propagates this information backwards to the internally generated tensors.

Fifth, *TorchLens* identifies and marks recurrent layers in the network. While the question of which layers should count as “recurrent” is a matter of definition, we stipulate that an operation is associated with a recurrent layer if it meets any of these conditions:

1. It contains trainable parameters and is executed more than once during the forward pass.
2. It is a parent or child of a pass of a layer meeting condition #1, and there is another operation of the same type adjacent to another pass of that layer (e.g., a ReLU operation following multiple passes of a recurrent convolutional layer). This condition is applied recursively (e.g., if the ReLU operation is in turn followed by a pooling layer in multiple passes), growing out a block of repeated layers surrounding the repeated-parameter layers, until no further layers can be added in this manner.
3. It is part of a sequence of operations that adjacently repeats, signifying a computational “loop”; corresponding operations in each pass of the loop are marked as passes through the same layer.

These conditions are illustrated in Figure 3.

To identify these repeated layers, we apply the following algorithm. First, candidate sets of “repeated layers” are identified in an initial first pass: these include layers in which the same trainable parameters are applied multiple times during the forward pass, and layers with no trainable parameters for which the same function with the same non-tensor arguments is applied multiple times during the forward pass (e.g., a ReLU operation, or scalar multiplication by a certain constant value). Second, the algorithm checks whether these candidate layers meet any of the above three criteria to count as “repeated”. To do this, the algorithm applies a procedure where it begins from any parentless layers (e.g., input layers or layers corresponding to an internally generated tensor) and steps forward one layer at a time, successively grouping together repeated sets of operations into blocks. For instance, for a given operation X (e.g., scalar multiplication by 10) that recurs multiple times, the algorithm identifies cases where that operation is also followed by recurring operations Y and Z (e.g., a ReLU operation and a pooling operation), identifying XYZ “blocks”. Third, after identifying these repeated blocks, it checks whether these blocks either contain a layer with trainable parameters, or repeat back to back (e.g., XYZXYZXYZ); if so, the corresponding operations in each block are marked as different passes of the “same” layer, completing the process.

Finally, all layer metadata is processed into its user-facing, human-readable form, and returned to the user.

### Validating the accuracy of saved activations

As noted above, it remains possible that unanticipated “edge case” scenarios may emerge for graphs with unexpected architectures, or that future updates to PyTorch may introduce elementary tensor functions that raise unique programming peculiarities. Thus, as a way of detecting such cases when they occur, and of ensuring the accuracy of any saved activations, *TorchLens* contains a built-in procedure for validating the accuracy of saved activations. This procedure works by executing the function (e.g., convolution) associated with a target layer on the saved values of the input layer(s) for that target layer, and verifying that the resulting output of this function call matches the saved tensor of that target layer. As a further check, it substitutes in random values instead of the saved values for the parent layers, and verifies that executing the function on these nonsense activations returns a tensor that differs from the saved tensor of the target layer (with additional logic that checks for “special case” like multiplying a tensor by zero, such that changing the input would not change the output). If the values of the target layer are saved correctly, then this procedure establishes that the parent layers of this target layer were also correctly saved. Since the final ground-truth outputs of the model are known, this procedure begins by using the model outputs as the “target” layer, and checking the parent layers to this target layer. If these parent layers pass the test, then they each become the new “target layer”, and the same procedure is applied to the parents of these layers. The procedure is then recursively applied until only layers with no parents (i.e., input layers and layers producing internally generated tensors) are left.

This approach has the benefit of requiring no manual troubleshooting, and is fully automatic. One drawback is that it logically requires saving the values of *all* intermediate tensors, which can impose high memory demands for large models.

### Compatibility

*TorchLens* is compatible with both CPU and GPU processing, and has been validated for PyTorch versions 1.9.0 and higher. Currently, it has not been tested in the case of parallel computing, although further updates to *TorchLens* are intended to address this.

## Discussion

*TorchLens* provides a method for not only automatically and exhaustively extracting the results of intermediate activations from arbitrary PyTorch DNN models, but also understanding the structure of these models via visualizations and metadata. We envision several possible use cases for *TorchLens*.

First, it can be used to streamline analysis pipelines for extracting and analyzing the internal representations encoded by DNNs for the purposes of better understanding their underlying computational principles and comparing these intermediate processing stages to those of the brain. Uniquely among existing approaches for this goal, *TorchLens* works automatically for arbitrary PyTorch models, exhaustively extracts the results of any desired operations without limitation, and provides metadata and visualizations to help characterize how a given layer fits into the broader architecture of the network. Since the ModelHistory object returned by *TorchLens* constitutes a full and easily queried representation of the model’s computational graph, it can also facilitate programmatic graph-theoretic analyses of a network, such as assessing how the similarity of the representations in two layers varies based on the nature of the intervening layers.

Second, *TorchLens* can be used as a prototyping, profiling, and visualization tool during model development. Since its visualization features provide an intuitive “picture of the code” for a model, it can provide a helpful visual aid when designing large or complicated models. Furthermore, the model metadata it provides includes information about the execution time of the functions in the model, and about the memory space used by the parameters and output tensors of all operations in the model, revealing runtime and storage bottlenecks. The automatic visualization functionality may also be useful when preparing presentations or publications describing a model.

Third, we envision TorchLens being pedagogically useful. For individuals who are first becoming acquainted with implementing deep learning models, the visualizations and information it provides can help in translating between model code and underlying concepts. Even for more seasoned practitioners, the ability to automatically generate a visual representation of a model can accelerate the process of understanding a new and complex model, which might otherwise require cumbersome inspection of a model’s code.

Fourth, *TorchLens* can be readily imported and used in other deep learning analysis pipelines. We note that while *TorchLens* can readily extract intermediate activations given a model and input, it does not have functionality for loading models or inputs, or for performing further analyses on the extracted activations. We make this design choice because we intend for *TorchLens* to be applicable in any of the many domains in which DNNs are used, making it infeasible to implement general functionality for loading models and analyzing activations. We note, however, that field-specific packages already exist for these purposes (e.g., Muttenthaler & Hebart, 2021; Nili et al., 2014).

While we intend for the current functionality of *TorchLens* to be largely self-contained, its ability to automatically extract a complete representation of the computational graph of a model does raise the potential for further applications built upon the same groundwork. One intriguing possibility is the ability to easily change or intervene upon an existing DNN model without directly changing the code, making it easier to apply causal inference methods (Pearl, 2009) to DNNs to evaluate causal claims about the effect of interventions, or to conduct counterfactual simulations. Since *TorchLens* stores the actual functions used in the forward pass of a model, the model can also be “re-run” from any intermediate stage, potentially with targeted changes introduced. For instance, if a user wishes to test how using a different nonlinearity function in a model changes a model’s internal representations, keeping all else the same, it would be simple to alter these nodes in the computational graph to use the new nonlinearity. Another example would be to “lesion” parts of a model by zeroing out particular parameters or values of a layer’s output tensor to examine the causal contribution of particular DNN units to a model’s behavior.

In summary, we hope that *TorchLens* will serve the role of removing arbitrary limitations to extracting and characterizing the results of internal DNN operations, streamlining both the scientific investigation and engineering of DNNs.

## Code and Data Availability

The open-source code for TorchLens is available on GitHub [here](https://github.com/johnmarktaylor91/torchlens). A gallery of visualizations for over 800 pretrained DNNs is available [here](https://drive.google.com/drive/u/0/folders/1BsM6WPf3eB79-CRNgZejMxjg38rN6VCb). A CoLab tutorial for TorchLens is available [here](https://colab.research.google.com/drive/1ORJLGZPifvdsVPFqq1LYT3t5hV560SoW?usp=sharing).

## Author Contributions

JT conceived of the project, JT and NK conceptualized *TorchLens* and prepared the manuscript, and JT implemented the package.

## Acknowledgements

We thank George Alvarez, Alfredo Canziani, Tal Golan, Colin Conwell, Jacob Prince, and members of the Visual Inference Lab at Columbia University for their helpful discussion and feedback during the development of *TorchLens*.

## Supplementary Information

**Supplementary Table 1:**
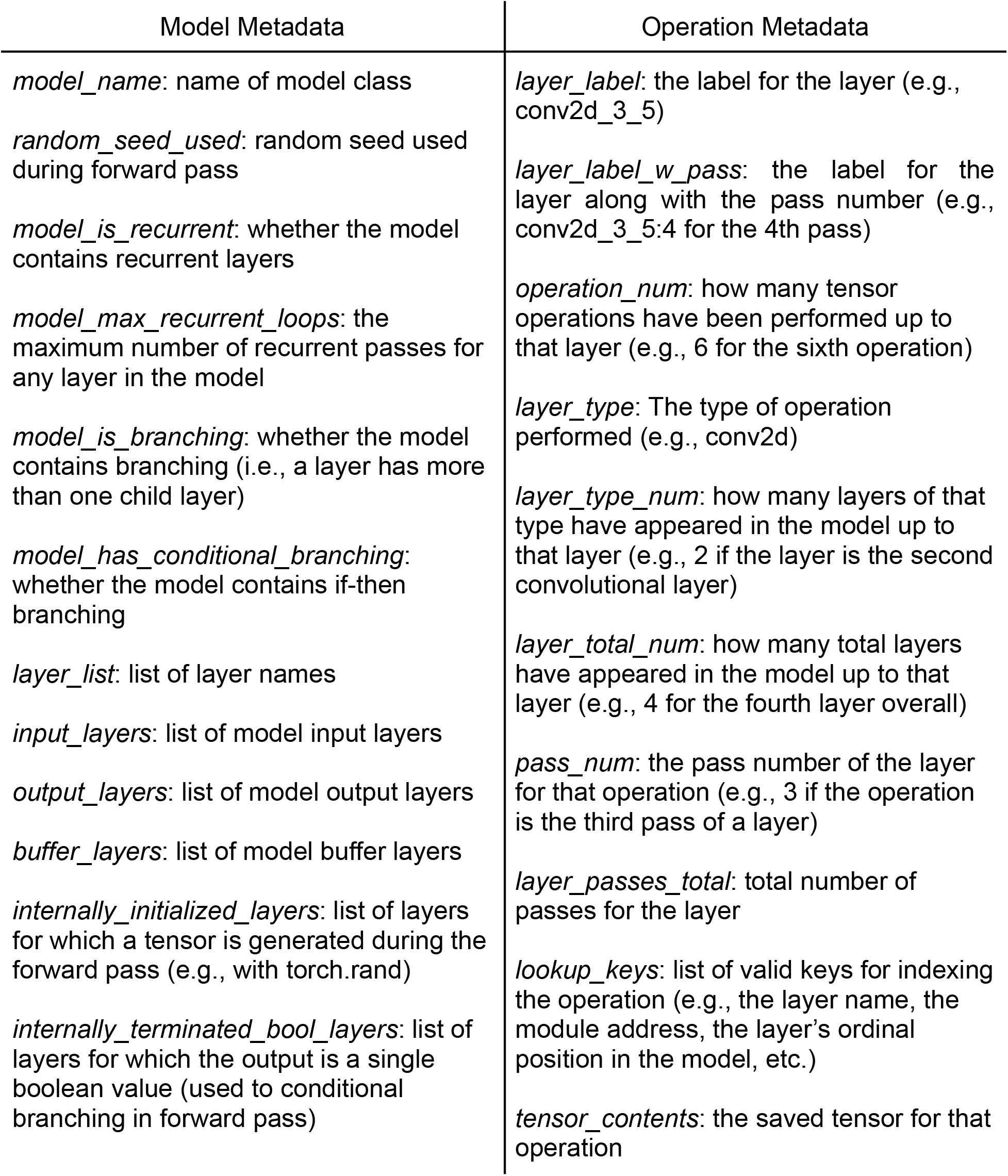
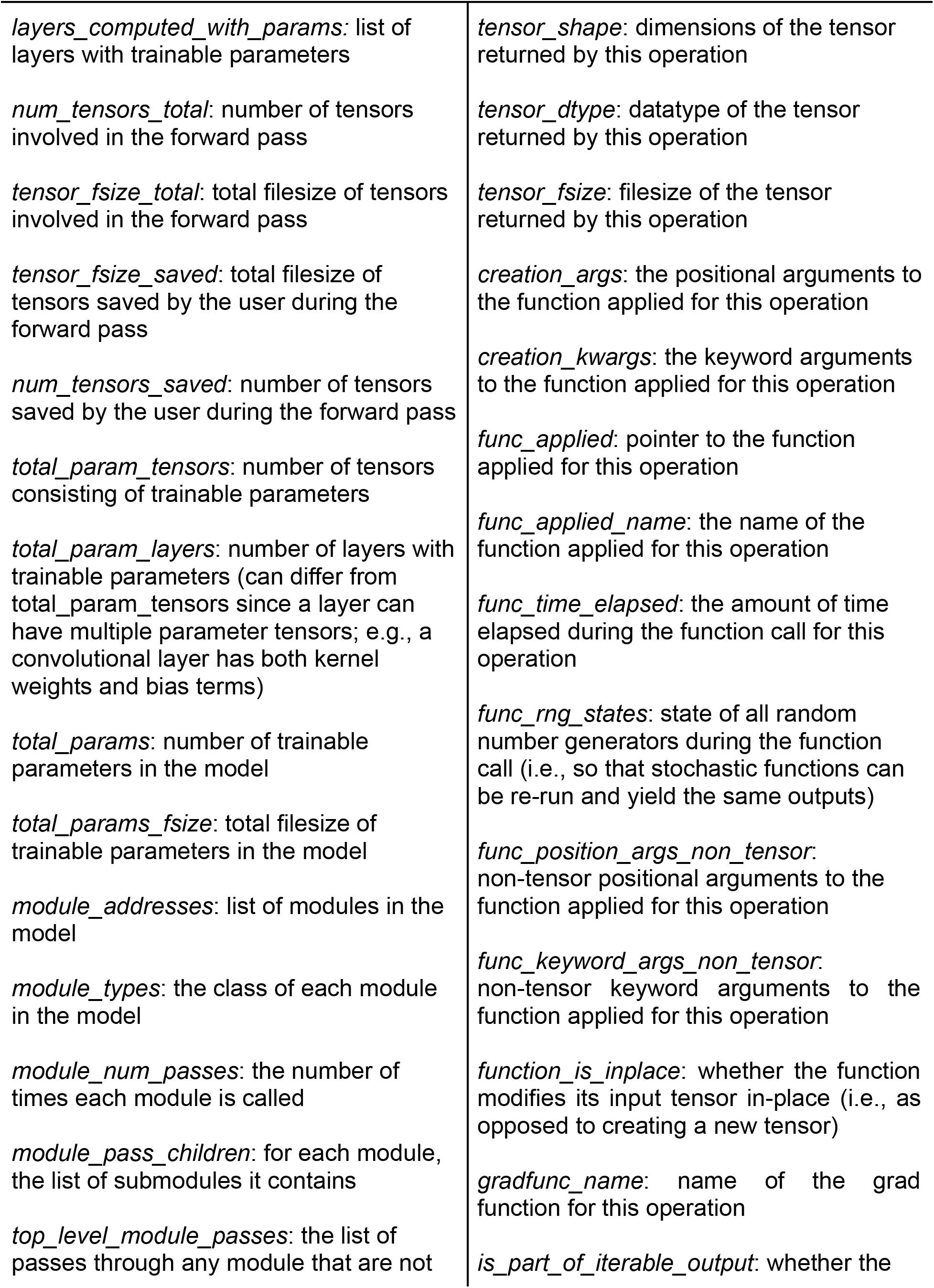
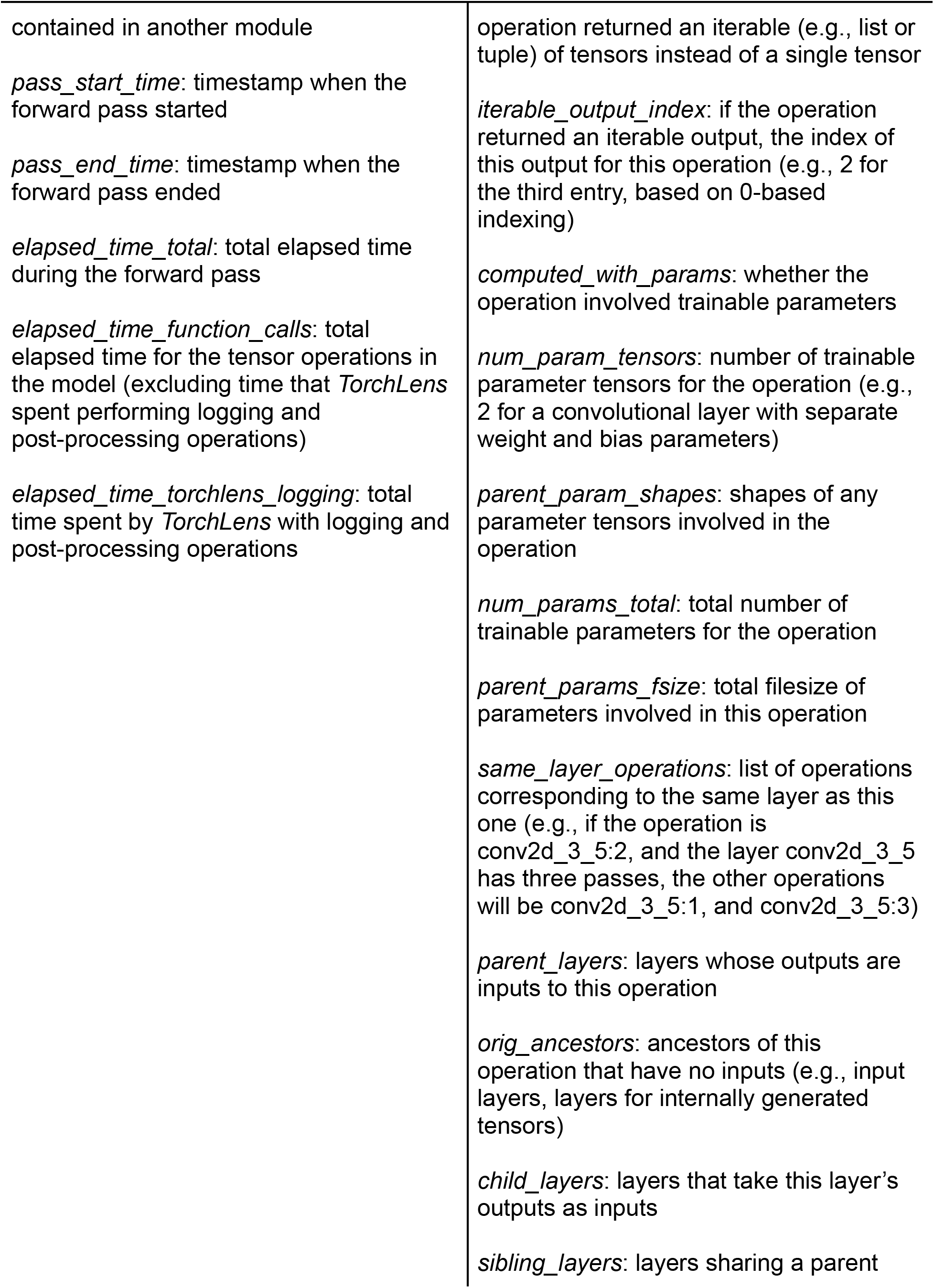
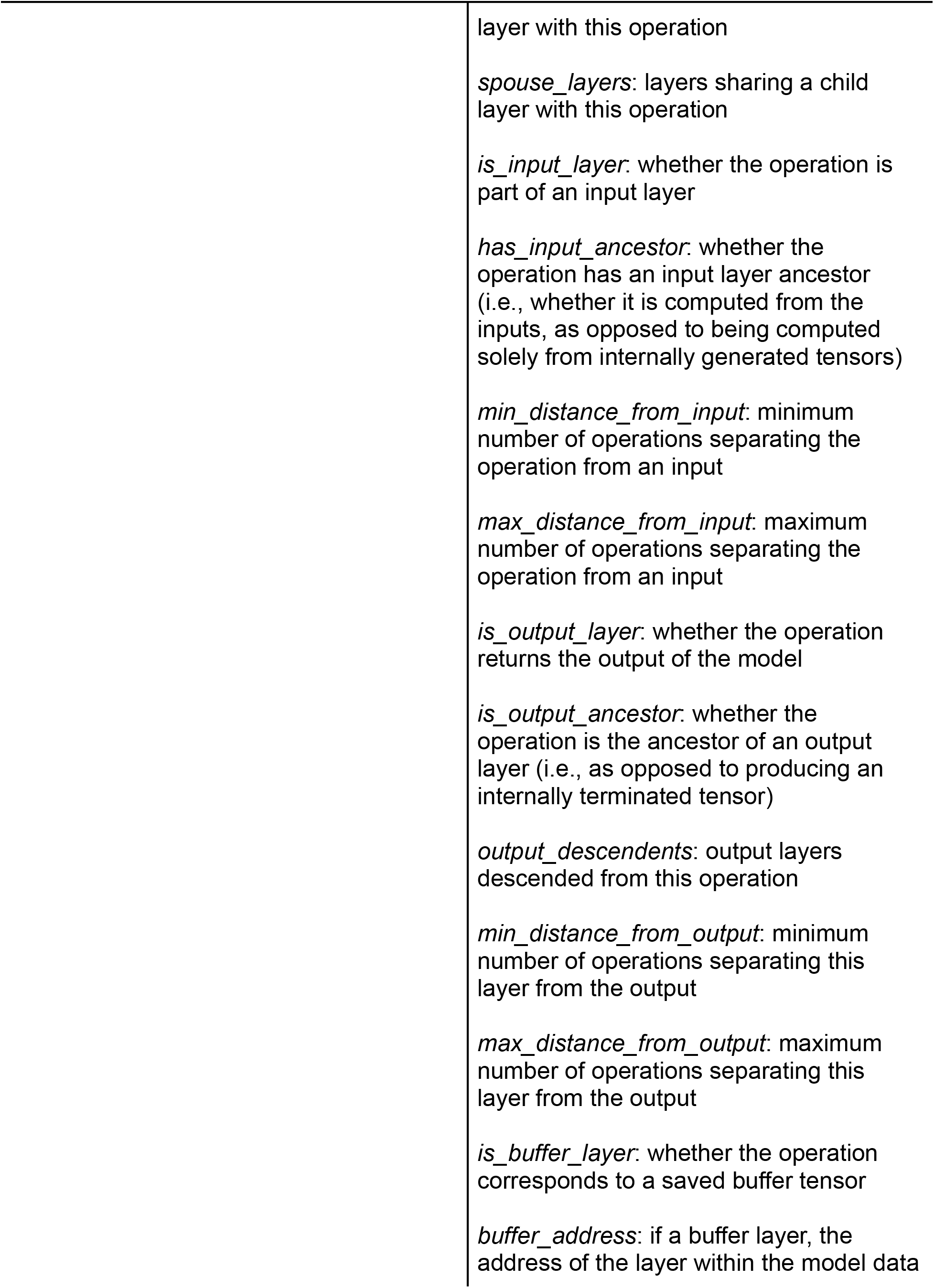
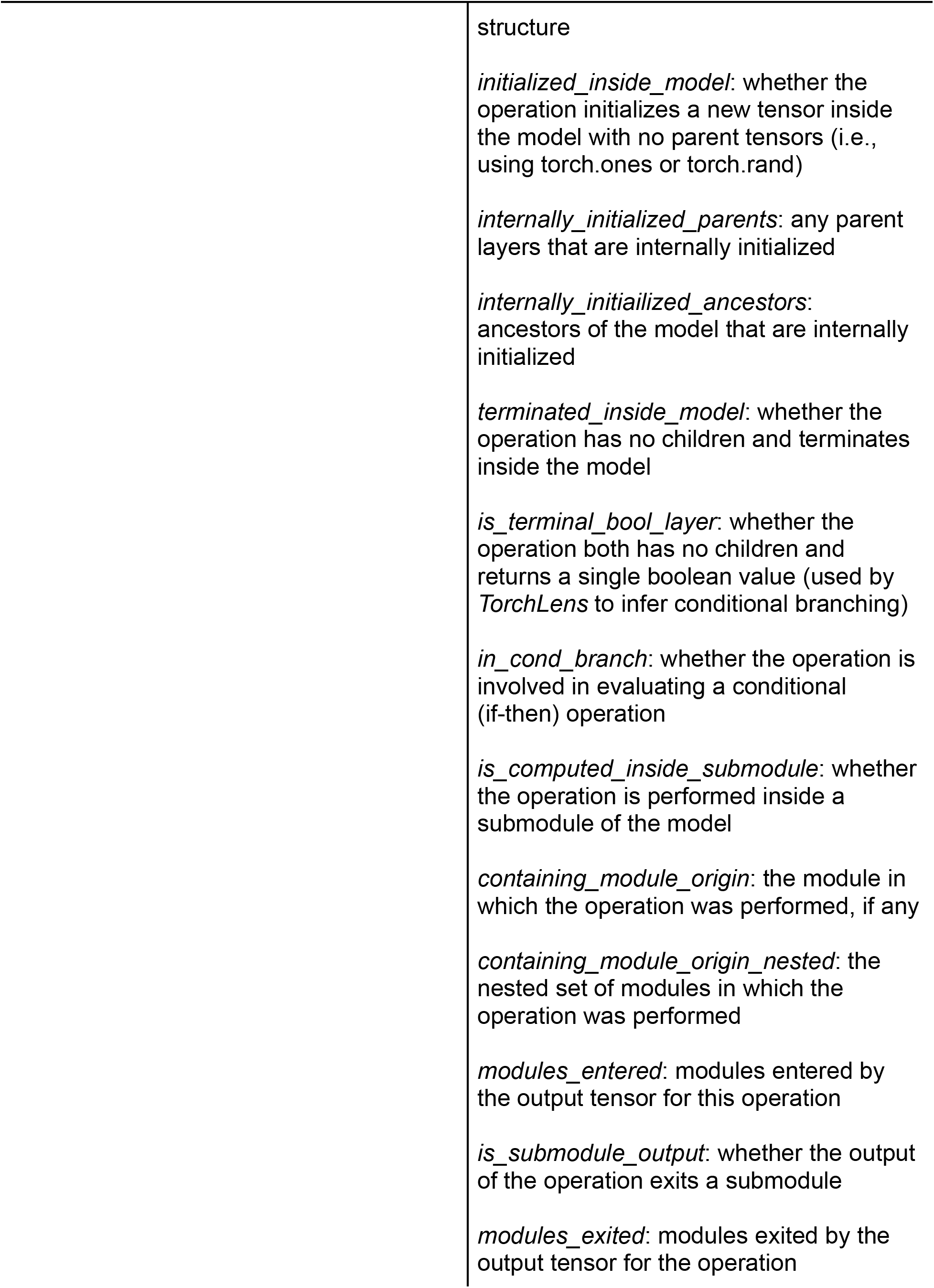
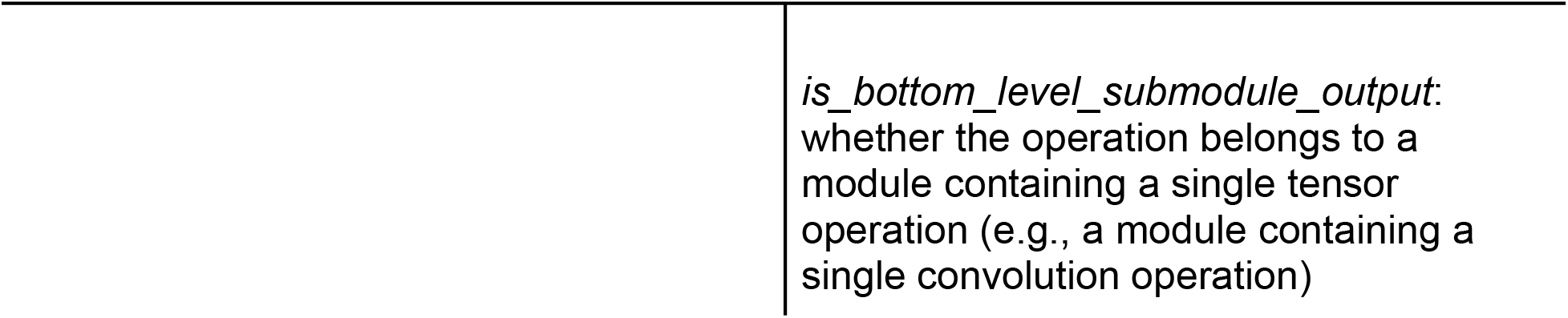
Model and layer metadata provided by *TorchLens*. Entries in the left column are attributes of the ModelHistory object returned by *TorchLens*; entries in the right are attributes of the log entry for each individual operation that can be fetched from ModelHistory.

## References

Devlin, J., Chang, M.-W., Lee, K., & Toutanova, K. (2019). BERT: Pre-training of Deep Bidirectional Transformers for Language Understanding (arXiv:1810.04805). arXiv. https://doi.org/10.48550/arXiv.1810.04805

Kearnes, S., McCloskey, K., Berndl, M., Pande, V., & Riley, P. (2016). Molecular graph convolutions: Moving beyond fingerprints. Journal of Computer-Aided Molecular Design, 30(8), 595–608. https://doi.org/10.1007/s10822-016-9938-8

Khaligh-Razavi, S.-M., & Kriegeskorte, N. (2014). Deep Supervised, but Not Unsupervised, Models May Explain IT Cortical Representation. PLOS Computational Biology, 10(11), e1003915. https://doi.org/10.1371/journal.pcbi.1003915

Kriegeskorte, N. (2015). Deep Neural Networks: A New Framework for Modeling Biological Vision and Brain Information Processing. Annual Review of Vision Science, 1(1), 417–446. https://doi.org/10.1146/annurev-vision-082114-035447

Kriegeskorte, N., Mur, M., & Bandettini, P. (2008). Representational Similarity Analysis – Connecting the Branches of Systems Neuroscience. Frontiers in Systems Neuroscience, 2.https://doi.org/10.3389/neuro.06.004.2008

Krizhevsky, A., Sutskever, I., & Hinton, G. E. (2012). ImageNet Classification with Deep Convolutional Neural Networks. In F. Pereira, C. J. C. Burges, L. Bottou, & K. Q. Weinberger (Eds.), Advances in Neural Information Processing Systems 25 (pp. 1097–1105). Curran Associates, Inc. http://papers.nips.cc/paper/4824-imagenet-classification-with-deep-convolutional-neural-networks.pdf

Kubilius, J., Schrimpf, M., Nayebi, A., Bear, D., Yamins, D. L. K., & DiCarlo, J. J. (2018). CORnet: Modeling the Neural Mechanisms of Core Object Recognition. BioRxiv, 408385. https://doi.org/10.1101/408385

Looks, M., Herreshoff, M., Hutchins, D., & Norvig, P. (2017). Deep Learning with Dynamic Computation Graphs (arXiv:1702.02181). arXiv. https://doi.org/10.48550/arXiv.1702.02181

Marcel, S., & Rodriguez, Y. (2010). Torchvision the machine-vision package of torch. Proceedings of the 18th ACM International Conference on Multimedia, 1485–1488. https://doi.org/10.1145/1873951.1874254

Mnih, V., Kavukcuoglu, K., Silver, D., Rusu, A. A., Veness, J., Bellemare, M. G., Graves, A., Riedmiller, M., Fidjeland, A. K., Ostrovski, G., Petersen, S., Beattie, C., Sadik, A., Antonoglou, I., King, H., Kumaran, D., Wierstra, D., Legg, S., & Hassabis, D. (2015). Human-level control through deep reinforcement learning. Nature, 518(7540), Article 7540. https://doi.org/10.1038/nature14236

Muttenthaler, L., & Hebart, M. N. (2021). THINGSvision: A Python Toolbox for Streamlining the Extraction of Activations From Deep Neural Networks. Frontiers in Neuroinformatics, 15. https://www.frontiersin.org/articles/10.3389/fninf.2021.679838

Nili, H., Wingfield, C., Walther, A., Su, L., Marslen-Wilson, W., & Kriegeskorte, N. (2014). A Toolbox for Representational Similarity Analysis. PLOS Computational Biology, 10(4), e1003553. https://doi.org/10.1371/journal.pcbi.1003553

Pearl, J. (2009). Causal inference in statistics: An overview. Statistics Surveys, 3(none), 96–146. https://doi.org/10.1214/09-SS057

Radford, A., Wu, J., Child, R., Luan, D., Amodei, D., & Sutskever, I. (2019). Language Models are Unsupervised Multitask Learners.

Schneider, F. (2022). Surgeon-pytorch. https://github.com/archinetai/surgeon-pytorch

Spoerer, C. J., Kietzmann, T. C., Mehrer, J., Charest, I., & Kriegeskorte, N. (2020). Recurrent neural networks can explain flexible trading of speed and accuracy in biological vision. PLOS Computational Biology, 16(10), e1008215. https://doi.org/10.1371/journal.pcbi.1008215

Xu, Y., & Vaziri-Pashkam, M. (2021). Limits to visual representational correspondence between convolutional neural networks and the human brain. Nature Communications, 12(1), Article 1. https://doi.org/10.1038/s41467-021-22244-7

Yamins, D. L. K., Hong, H., Cadieu, C. F., Solomon, E. A., Seibert, D., & DiCarlo, J. J. (2014). Performance-optimized hierarchical models predict neural responses in higher visual cortex. Proceedings of the National Academy of Sciences, 111(23), 8619–8624. https://doi.org/10.1073/pnas.1403112111

Yu, Y., Abadi, M., Barham, P., Brevdo, E., Burrows, M., Davis, A., Dean, J., Ghemawat, S., Harley, T., Hawkins, P., Isard, M., Kudlur, M., Monga, R., Murray, D., & Zheng, X. (2018). Dynamic control flow in large-scale machine learning. Proceedings of the Thirteenth EuroSys Conference, 1–15. https://doi.org/10.1145/3190508.3190551

